# Platelets induce cell apoptosis of cardiac cells via FasL after acute myocardial infarction

**DOI:** 10.1101/2024.03.20.585918

**Authors:** Kim J. Krott, Friedrich Reusswig, Matthias Dille, Evelyn Krüger, Simone Gorressen, Saoussen Karray, Amin Polzin, Malte Kelm, Jens-W. Fischer, Margitta Elvers

**Affiliations:** Department of Vascular and Endovascular Surgery, Experimental Vascular Medicine, Heinrich-Heine-University Medical Center, Düsseldorf, Germany; Institute for Pharmacology and Clinical Pharmacology, Heinrich-Heine University, Düsseldorf, Germany; Université de Paris, INSERM U976 F-7510 Paris, France; Department of Cardiology, Pulmonology and Angiology, Heinrich-Heine University Medical Center, Düsseldorf, Germany

**Keywords:** Platelets, myocardial infarction, apoptosis, cardiomyocytes, Fas ligand

## Abstract

Acute myocardial infarction (AMI) is one of the leading causes of death worldwide. Cell apoptosis in the myocardium plays an important role in ischemia and reperfusion (I/R) injury, leading to cardiac damage and dysfunction. Platelets are major players of hemostasis and play a crucial role in vessel occlusion, inflammation and cardiac remodeling after I/R. Here, we studied the impact of platelets on cell apoptosis in the myocardium using a close-chest mouse model of AMI. We found caspase-3 positive resident cardiac cells while leukocytes were negative for caspase-3. Using two different mouse models of thrombocytopenia, we detected a significant reduction of caspase-3 positive cells in the infarct border zone after I/R injury. Further, we identified platelet FasL to induce cell apoptosis via the extrinsic pathway of Fas receptor activation of target cells. Mechanistically, hypoxia triggers platelet adhesion to FasR suggesting that platelet induced apoptosis is elevated after I/R. Platelet-specific FasL knock-out mice showed reduced Bax and BcL-2 expression suggesting that platelets modulate the intrinsic and the extrinsic pathway of apoptosis leading to reduced infarct size after myocardial I/R injury. Therefore, platelet induced cardiac damage needs to be taken into account while optimizing antithrombotic/antiplatelet strategies for patients with AMI.

## 1. Introduction

Ischemia and reperfusion (I/R) injury induces lethal injury to cardiomyocytes during acute myocardial infarction. There is growing evidence for cell apoptosis to play a major role in myocardial I/R injury. Extrinsic and intrinsic pathways of cell death might play a role in I/R induced cell apoptosis including mitochondria and cell death receptor FasR (1, 2). In addition, other mechanisms such as necrosis and autophagy might play a role in cardiac damage after acute myocardial infarction (AMI) (3-5). Thus, inhibitors of these pathways have been shown to be cardioprotective (6).

Platelets are anuclear blood cells with a strong impact on hemostasis and thrombosis. At sites of vascular damage, platelets adhere and accumulate to restore tissue integrity and to avoid massive blood loss. However, uncontrolled platelet activation at sites of atherosclerotic plaque rupture is an important factor of ischemic events as observed in patients with AMI (7). According to the World Health Organization (WHO) ischemic heart diseases are still the global leading cause of death. Obstructed coronary vessels are recanalized by percutaneous coronary intervention (PCI) that often leads to I/R injury, which can cause additional cardiac damage in patients with AMI (8, 9).

Different knock-out or blocking strategies in experimental mice and rats as well as human studies revealed a significant contribution of platelets to coronary thrombosis, acute inflammation and cardiac damage after AMI (10-17). Recently, we have shown that elevated platelet activation after AMI contributes to I/R injury in patients and in mice (14). Enhanced platelet activation leads to migration of platelets into the infarct border zone in the left ventricle (LV) where platelets might interact with resident cardiac cells. This allows platelets to modulate cardiac remodeling beside their role in acute inflammation by interacting with fibroblasts after AMI.

However, a role for platelets in cell apoptosis has been described recently (18). Activated platelets induce cell apoptosis in murine neuronal cells, human neuroblastoma cells, and mouse embryonic fibroblasts by the exposure of FasL at the platelet surface. Consequently, platelet depletion resulted in reduced apoptosis in an experimental mouse model of stroke (18). However, nothing is known about the impact of platelets in cell death after AMI.

In this study, we analyzed the capacity of platelets to substantially impact cell survival in the infarct border zone after myocardial ischemia and reperfusion. Our results provide strong evidence for platelets to induce apoptosis of resident cardiac cells in the LV via the intrinsic and the extrinsic pathway as shown by mitochondrial and FasL-induced mechanisms.

## 2. Materials and Methods

### 2.1 Animals

The animal experiments were conducted following the guidelines established by the European Parliament for the ethical use of live animals in scientific research and in compliance with the German legislation concerning animal welfare. Approval for the experimental protocol was obtained from both the Heinrich Heine University Animal Care Committee and the district government of North-Rhine-Westphalia (LANUV; NRW), under Permit Numbers 84-02.04.2015.A558, 81-02.04.2019.A270, and 84-02.04.2017.A440. Wildtype (C57BL/6J) mice were purchased from Janvier (Le Genest-Saint-Isle, France).

Wildtype mice were used for induction of thrombocytopenia. Platelet depletion was initiated by administering a GPIba antibody (No #R300, Emfret, Eibelstadt, Germany) or the corresponding IgG-control antibody (Order No #C301, Emfret, Eibelstadt, Germany) 24 hours before ligating the left anterior descending coronary artery (LAD). Platelet depletion, general blood cell counts, and isolated platelet counts at various time points after ischemia/reperfusion (I/R) were monitored using the automated hematology analyzer Sysmex (Sysmex Corporation, Kobe, Japan) as already described.

Furthermore, a C57BL/6-FasL-floxed mouse strain (Inserm, Paris) was crossbred with the PF4-Cre recombinase-expressing mouse strain (C57BL/6-Tg(PF4-cre)Q3Rsko/J; The Jackson Laboratory, USA) to facilitate the specific knock-out of genes in megakaryocytes and platelets. Wildtype littermates corresponding to the strains were bred from parental pairs and genotyped through PCR analysis.

For surgical procedures, mice were anesthetized via intraperitoneal injection with Ketamine (100 mg/mL, Ketaset, Zoetis, Malakoff, France) and Xylazine (20 mg/mL, WDT, Ulft, Netherlands). Euthanasia was performed by cervical dislocation.

### 2.2 Experimental model of acute myocardial infarction (AMI) and reperfusion in mice

In order to mitigate surgical trauma and subsequent inflammatory reactions resulting from intervention and antibody injection following ischemia/reperfusion (I/R), a closed-chest model of reperfused myocardial infarction was employed (19). Male mice aged 10 to 12 weeks were subjected to anesthesia via intraperitoneal injection of a solution containing Ketamine (90 mg/kg body weight) and Xylazine (10 mg/kg body weight). Upon successful induction of anesthesia, the left anterior descending artery (LAD) was ligated for 60 minutes to induce myocardial infarction (MI) three days post instrumentation. Coronary occlusion was achieved by gradually tightening the applied suture until ST-elevation appeared on the electrocardiogram (ECG). Reperfusion was confirmed by the resolution of ST-elevation. After 24 hours of reperfusion, hearts were excised and stained with TTC/Evans Blue solution to delineate the damaged area, distinguishing between the area at risk (ischemic area) and the infarcted area. The ratios of these distinct areas were digitally quantified using video planimetry. Left ventricular function post-MI was assessed via echocardiography at various time points after I/R (1d, 5d and 21d), utilizing a Vevo 2100 ultrasound machine (VisualSonics Inc., Bothell, USA) to measure parameters including ejection fraction (%), cardiac output (mL/min), fractional shortening (%), and stroke volume (μL), with corresponding software.

### 2.3 Experiments with human blood and study populations

The experiments involving human blood were reviewed and approved by the Ethics Committee of Heinrich-Heine-University, adhering to the principles of the Declaration of Helsinki. Written consent was obtained from all participants before their involvement in the study. In general, fresh citrate-anticoagulated blood (105 mM Na_3_-citrate, Becton Dickinson, New Jersey, USA) was collected from healthy volunteers aged 18 to 70 years in the Blood Donation Center at the University Hospital Düsseldorf for experiments under hypoxic conditions. To measure sFasL in Plasma of STEMI patient, blood samples were collected from 30 STEMI and 18 CCS patients using 2.7 mL 0.1 M sodium citrate tubes (BD Vacutainer®, Becton Dickinson, New Jersey, USA) and promptly centrifuged to isolate plasma samples for sFasL analysis.

For the STEMI patient cohort, eligible patients admitted to the University Hospital Düsseldorf were screened for inclusion and exclusion criteria. Inclusion criteria comprised ST-elevation myocardial infarction (STEMI) or chronic coronary syndrome (CCS) within existing coronary artery disease (CAD). Exclusion criteria included a palliative situation (life expectancy less than 8 weeks), pregnancy, cognitive impairment, dementia, severe chronic kidney disease (stage 3b-5), severe liver dysfunction, and platelet disorders. To mitigate potential biases, patients’ characteristics were compared between groups.

### 2.4 Human platelet preparation

Human blood fresh citrate-anticoagulated blood (105 mM Na_3_-citrate, BD-Vacutainer^®^; Becton, Dickinson and company) was collected and centrifuged at 200 *g* for 10 min without brake. The upper phase consisting of the platelet-rich plasma (PRP) was carefully transferred into phosphate buffered saline (PBS) pH 6.5 containing apyrase (2.5 U/mL) and 1 μM PGE_1_. The tubes were centrifuged at 1000 *g* for 6 min and the obtained pellet was resuspended in tyrode’s buffer (137 mM NaCl, 2.8 mM KCl, 12 mM NaHCO_3_, 0.4 mM NaH_2_PO_4_, 5.5 mM glucose, pH 6.5). The final cell count was measured by a hematology analyzer (Sysmex KX-21N, Norderstedt, Germany) and adjusted according to the requirements of the experiment.

### 2.5 Platelet adhesion to immobilized Fas receptor protein

Adhesion experiments on immobilized FasR protein were performed with isolated human platelets. Isolated platelets were pre-incubated for 30 min in HEPES buffer (10 mM HEPES; 150 mM NaCl; 10 mM D-Glucose; 1 mM MgCl_2;_ 5 mM KCl; pH 7.4) under normoxic (21% O_2_) or hypoxic (2% O_2_) conditions in the Whitley H35 HEPA Hypoxystation. Glass coverslips (24 × 60 mm) were coated with human recombinant Fas protein (50 μg/mL, Sino Biological) at a defined area (10 × 10 mm); incubated in humidity chambers at 4 °C overnight. Coverslips were blocked with 1% BSA for at least 1 h at RT. Resting or ADP (10 μM)-stimulated platelets (4 × 10^3^/μL) were pretreated with the inhibitor hDcR3 (10 μg/mL, recombinant human decoy receptor 3 protein, R&D Systems) for 10 min and then allowed to adhere to the coated coverslips for 60 min at RT under normoxic or hypoxic conditions. IgG-Fc (10 μg/mL, R&D Systems) peptide served as control. Afterwards, cover slips were rinsed two times with PBS to wash off unbound platelets. Adherent cells were fixated immediately with 4% paraformaldehyde (PFA) and covered with the mounting medium (Aquatex^®^). Five DIC images from different areas were taken (platelets: 400-fold total magnification; CHO cells: 100-fold total magnification; Axio Observer.D1, Carl Zeiss). The total number of adherent platelets was counted using ImageJ-win64 software.

### 2.6 Flow cytometric analysis of blood cells

To analyze Fas-Ligand (FasL) and PS exposure on the platelet surface, the expression was measured via fluorescence-based flow cytometry. CD42b was used as platelet specific marker in all experiments.

For detection of the PS exposure Cy™5 AnnexinV (BD Biosciences) staining was conducted while binding buffer (10mM Hepes, 140 mM NaCl, 2.5 mM CaCl2, pH 7.4) was used instead of PBS. FasL exposition and PS exposure was measured on platelets (CD42b^+^ population) in washed whole blood 24 hours after MI. Samples were incubated Annexin Cy^TM5^ or PE-labeled FasL antibody (CD178-PE, NOK1, Biolegend) for 15 min at RT. Reactions were stopped by adding of 500μL PBS or AnnexinV binding puffer, respectively, and measured at a FASCalibur (BD). MFI of FasL was measured in the platelet population, representing the FasL exposition on platelets.

For PS exposure and FasL externalization analysis under normoxic (21% O_2_) or hypoxic (2% O_2_) conditions, isolated human platelets were pre-incubated for 30 min under these conditions in HEPES buffer (10 mM HEPES, 150 mM NaCl, 10 mM D-Glucose, 1 mM MgCl_2_, 5 mM KCl, pH 7.4) in the Whitley H35 HEPA Hypoxystation. Resting or CRP-activated platelets were labeled and analyzed as described before.

### 2.7 Enzyme-Linked Immunosorbent Assay (ELISA)

For quantification of soluble FasL in plasma of mice, heparinized blood was centrifuged 10 min for 650 g to collect the plasma. The amount of soluble FasL was measured by an ELISA Kit from My Biosource, San Diego, USA, following the manufacturer′s protocol.

For detection of sFasL in STEMI Patients plasma was collected in EDTA tubes. Samples were centrifuged at 1,000 g for 30 minutes before proceeding to ELISA (Human FAS Ligand ELISA Kit - CD95L –Abcam-ab100515).

### 2.8 Immunohistochemistry of cardiac sections

At different time points after ischemia/reperfusion hearts were flushed with cold heparin solution (20 U/mL, Roche, Basel, Switzerland), removed, paraffin embedded and cut into 5 μm sections using an automatic microtome (Microm HM355, Thermo Fisher Scientific). Prior to staining, all sections were deparaffinised and hydrated. For antigen unmasking, the tissue sections were heated at 300 W in citrate buffer (pH 6.0) for 10 min. For fluorescence staining of apoptotic cells in the LV, the following antibodies were used: Cleaved caspase-3 (Casp-3, #9661, Cell Signaling, 1:50), vimentin (#PA5-142829, Invitrogen, 1:50), troponin (#MA5-12960, Invitrogen, 1:50) and CD45 (#55307, Cell Signalling, 1:100). The heart tissue sections were blocked with DPBS (containing 0.3% Triton x-100 and 5% goat serum) for 1h at RT. After overnight incubation with the respective primary antibodies at 4 °C, the sections were stained with a highly cross-adsorbed Alexa Fluor™ Plus 555 conjugated secondary antibody (#A32794, Invitrogen™, 1:200) and Alexa Fluor™ 647 conjugated secondary antibody (#A21247, Invitrogen™, 1:200) for 1h at RT. In control experiments, the first antibodies were omitted and the sections were only stained with secondary antibodies. For analyzing apoptosis in thrombocytopenic or FasL knock-out mice after myocardial infarction, paraffin-embedded heart section were immune-stained with cleaved Caspase-3 as primary antibody, followed by labeling with streptavidin biotin/horseradish peroxidase (LSAB2 System HRP, Dako, Santa Clara, USA) and Diaminobenzidine (DAB) - Chromogen (Dako, Santa Clara, USA), performed as standard protocol. For visualising nuclei, all tissue sections were stained with DAPI (4’, 6-Diamidine-2’-phenylindole dihydrochloride, #10236276001, Roche, 1:3000). Images were taken with a confocal microscope from Zeiss (LSM 880 Airyscan).

### 2.9 Quantitative Real-Time polymerase chain reaction (qRT-PCR)

To evaluate the endogenous expression levels of Bax and Bcl-xl, isolated total RNA from the LV 24 hours post-myocardial infarction was utilized. LVs were separated and homogenized in Trizol using the Precellys tissue homogenizer (Precellys 24-Dual Homogenizer, Bertin, Frankfurt am Main, Germany) for 30 seconds at 5000 rpm. The supernatant was collected, chloroform was added, and RNA isolation was carried out using an RNAeasy Mini Kit (Qiagen), following the manufacturer’s instructions. Following reverse transcription, quantitative PCR amplification was conducted using the following oligonucleotide primers:

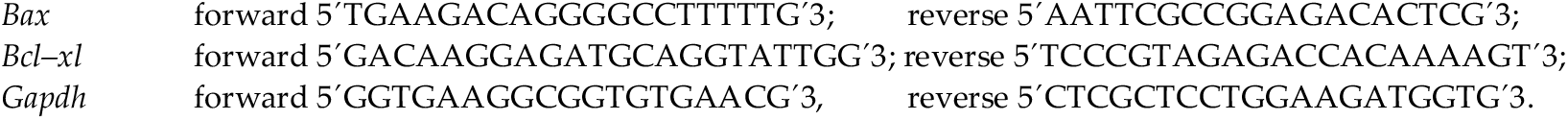

Quantitative real-time PCR was performed using Fast Sybr Green Master Mix (Thermo Fischer Scientific, Waltham, USA) according to a standard protocol. The expression levels of *Bax* or *Bcl-xl* were normalized to glyceraldehyde-3-phosphate dehydrogenase (Gapdh) RNA expression levels from naïve control mice.

### 2.10 Statistical analysis

The experiments were conducted a minimum of three times, with ‘n’ representing each individual animal. Results are expressed as means ± SEM. Statistical analysis was conducted using GraphPad Prism 10.0 software (Graphpad Software, Inc, San Diego, USA). Statistical tests included t-tests, one-way, or two-way ANOVA, with significance set at P < 0.05. In all figures, significance levels were denoted as *P < 0.05, **P < 0.01, and ***P < 0.001.

## 3. Results

### 3.1 Ischemia and reperfusion induced cell apoptosis of resident cardiac cells in the infarct border zone after AMI in mice

After acute myocardial infarction, ischemia and reperfusion injury leads to cell apoptosis in the myocardium (20). To identify cells in the myocardium that are affected by cell apoptosis, we first performed a histological analysis using different cellular markers to stain cardiomyocytes (troponin), cardiac fibroblasts (vimentin) and leukocytes (CD45) that have been migrated into the infarct border zone. As shown in figure 1, we detected caspase-3 positive cardiomyocytes (Fig. 1A-B) and caspase-3 positive fibroblasts (Fig. 1C-D) in the infarct border zone of mice that underwent acute myocardial infarction using cardiac sections of the LV at day 5 post I/R. However, the analysis of CD45 positive leukocytes revealed that these cells are not affected by apoptosis (Fig. 1E-F). Thus, only resident cardiac cells but not migrated inflammatory cells showed signs of apoptosis as detected by caspase-3 positive cardiomyocytes and fibroblasts.

**Figure 1.**
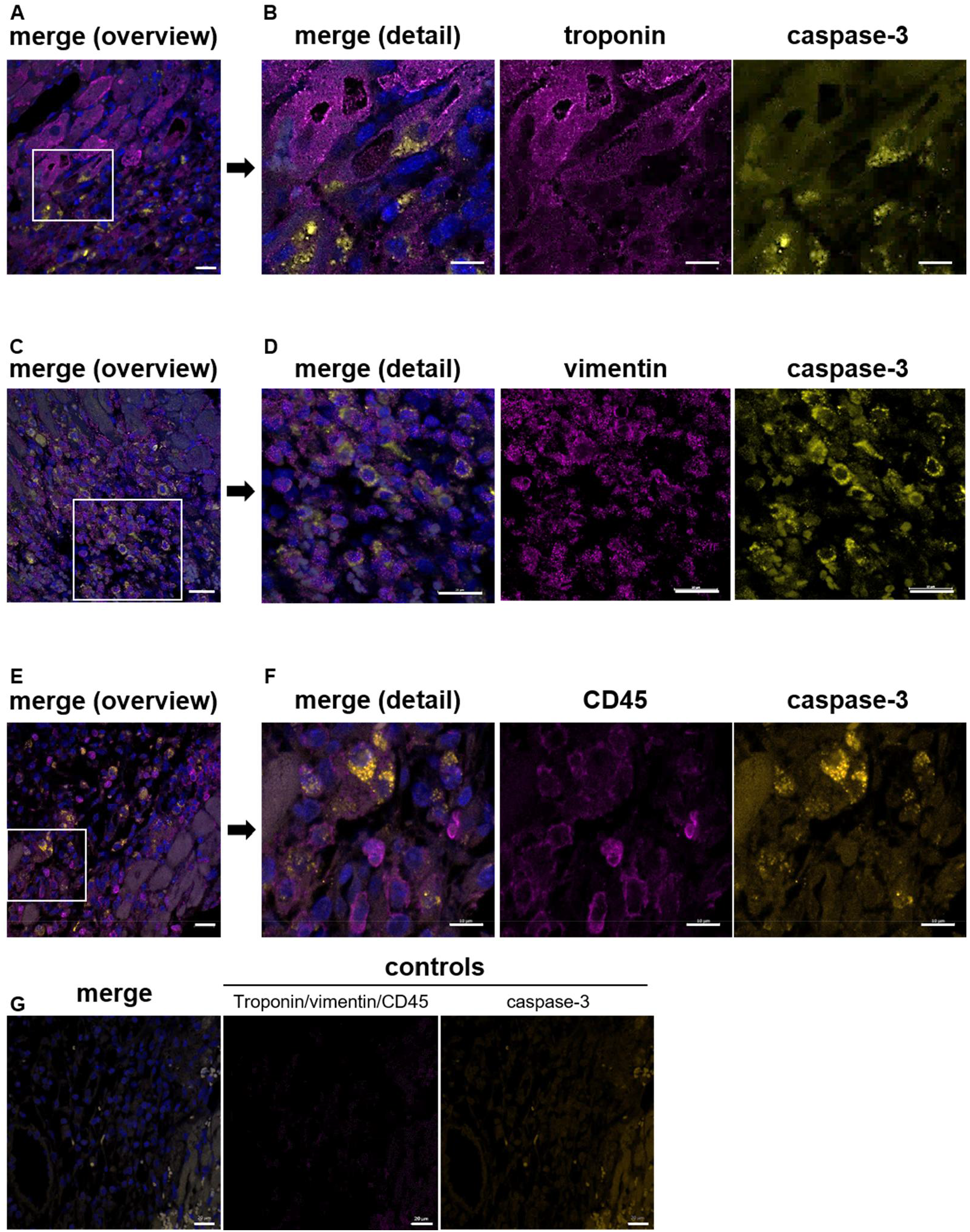
Ischemia and reperfusion injury leads to apoptosis of resident cardiac cells in the infarct border zone 5 days after AMI in mice. **(A-B)** Representative images of cardiac sections from the infarct border zone 5 days after I/R stained for troponin as marker for cardiomyocytes and caspase-3, **(A)** overview, **(B)** section from A to show details of caspase-3 positive cardiomyocytes; n = 4. **(C-D)** Caspase-3 positive cardiac fibroblasts in the infarct border zone were detected 5 days post AMI. **(C)** Overview, **(D)** details of vimentin stained fibroblasts; n = 3-5. **(E-F)** CD45 stained leukocytes show no signs of apoptosis using cardiac sections of the infarct border zone 24 h post I/R. **(E)** Overview and details **(F)** of CD45 stained leukocytes and caspase-3 positive cells were shown; n = 4-6. **(G)** Controls of troponin/vimentin/CD45 and caspase-3 staining were shown using PBS and the appropriate secondary antibody (the same secondary antibody was used for troponin, vimentin and CD45). In all samples, DAPI was used to stain nuclei in cardiac sections. Scale bar, (**A, C, F** and **G)** = 20 μm, (**B** and **D)** = 10 μM, **(E)** = 30 μM.

### 3.2 Reduced cardiac cell apoptosis in thrombocytopenic mice after acute myocardial infarction

Cell apoptosis plays a crucial role in LV remodeling after AMI (21). Several extracellular signals regulate apoptosis in cardiomyocytes, which ends up in the activation of caspase-3 (22). To analyze if platelets play a role in cell apoptosis in the infarct border zone after I/R, we analyzed platelet depleted and MPL knock-out mice that suffer from thrombocytopenia. First, we determined the number of caspase-3 positive cells in the infarct border zone of thrombocytopenic mice by histologic analysis of infarcted heart sections. As shown in figure 2A, a significant reduction of apoptotic cells has been observed in mice with almost no platelets (platelet depleted mice) and in MPL knock-out mice with platelet counts of ∼10% (14). To investigate which mechanisms play a role in platelet induced cell apoptosis post AMI, we first investigated mitochondria-dependent apoptosis via the intrinsic pathway. A lower expression of the apoptosis associated gene *Bax* was found in the LV of platelet depleted mice after 24h of reperfusion (Fig. 2B) while non-depleted control mice showed an enhanced *Bax* gene expression due to ischemia and reperfusion injury (Fig. 2B). In line with enhanced *Bax* expression, an increase in gene expression of the anti-apoptotic *Bcl*-2 in the LV was observed in mice where platelets have been depleted after AMI while no differences were observed in IgG control mice (Fig. 2C). In contrast, chronic thrombocytopenic MPL knock-out mice displayed no differences in *Bax* and *Bcl*-2 gene expression in the LV 24h after ischemia and reperfusion injury (Fig. 2B-C).

**Figure 2.**
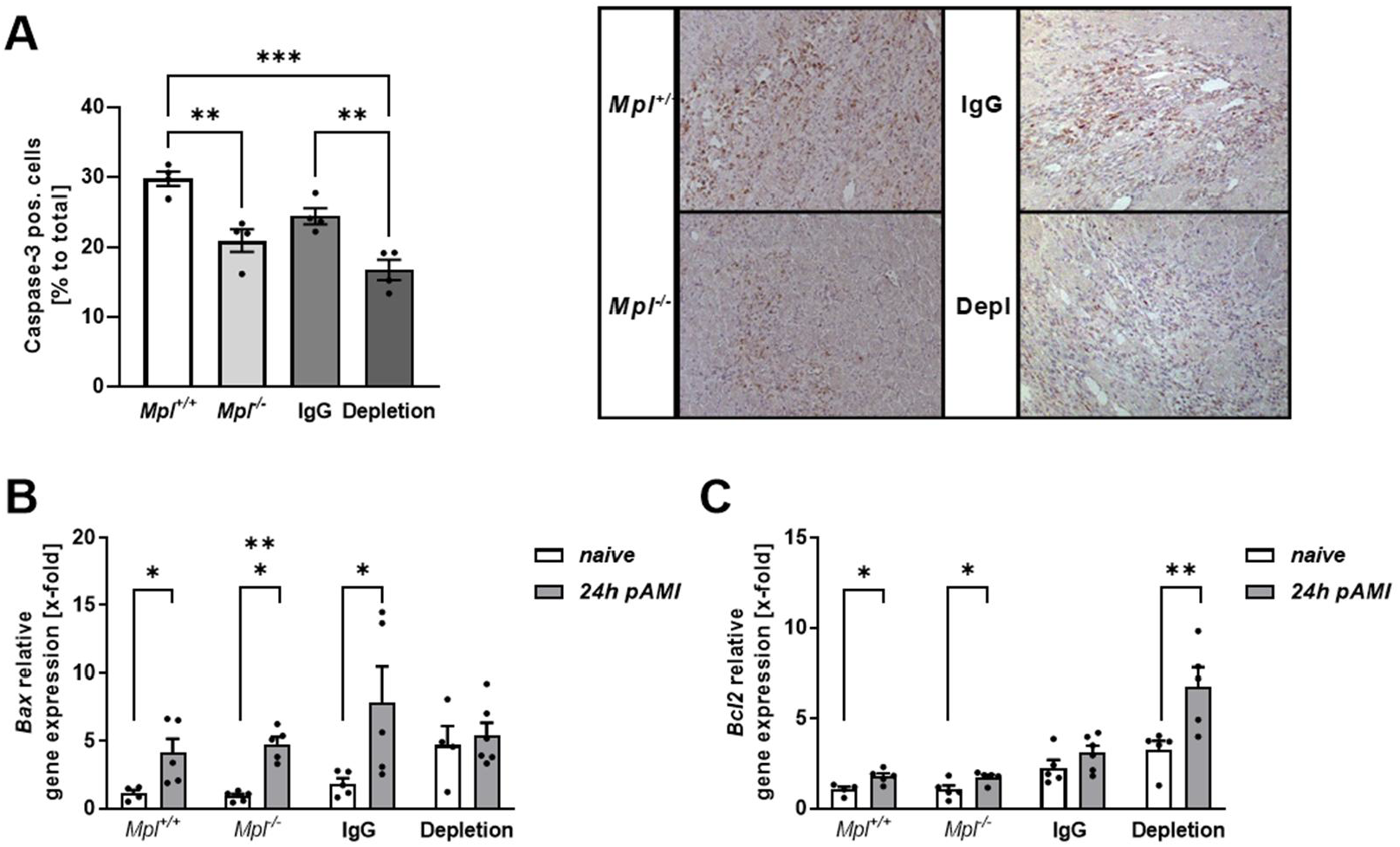
Reduced cardiac cell apoptosis in thrombocytopenic mice after acute myocardial infarction. **(A)** Quantification (left panel) and representative images (right panel) of caspase-3 positive cells in the infarct border zone of platelet depleted and Mpl deficient mice compared to controls 5 days after AMI; n = 4. **(B-C)** mRNA analysis of pro-apoptotic *Bax* **(B)** and anti-apoptotic *Bcl-2* **(C)** genes in the LV (left ventricle) of thrombocyopenic naive mice and 24h post AMI compared to respective control mice; n = 4-6 **(B);** n = 4-6 **(C)**. Scale bar = 50 μm. Data are presented as means ± SEM. Statistical analyses were done by **(A)** Two-Way ANOVA followed by Sidak post hoc and **(B and C)** multiple t-test. *p < 0.05, **p < 0.01, ***p < 0.001.

### 3.3 Increased adhesion of platelets to immobilized Fas receptor under hypoxic conditions

Beside cell apoptosis via the intrinsic pathway, cells can undergo apoptosis by the extrinsic pathway via TNF or FasL (23). The FasL-FasR plays an important role in the extrinsic pathway of cell apoptosis (24). Shedding of the FasL to its soluble form (sFasL) is associated with the loss of its apoptotic function because sFasL blocks the activity of FasR (25). Platelets are known to expose FasL at the membrane that is associated with platelet-RBC interaction (26) and platelet-induced cell apoptosis (18). Therefore, we next analyzed the impact of platelet FasL binding to FasR under hypoxic conditions. As shown in figure 3A, platelets adhere to immobilized recombinant FasR under physiological oxygen levels. However, hypoxic conditions induced increased adhesion of platelets to immobilized FasR.

**Figure 3.**
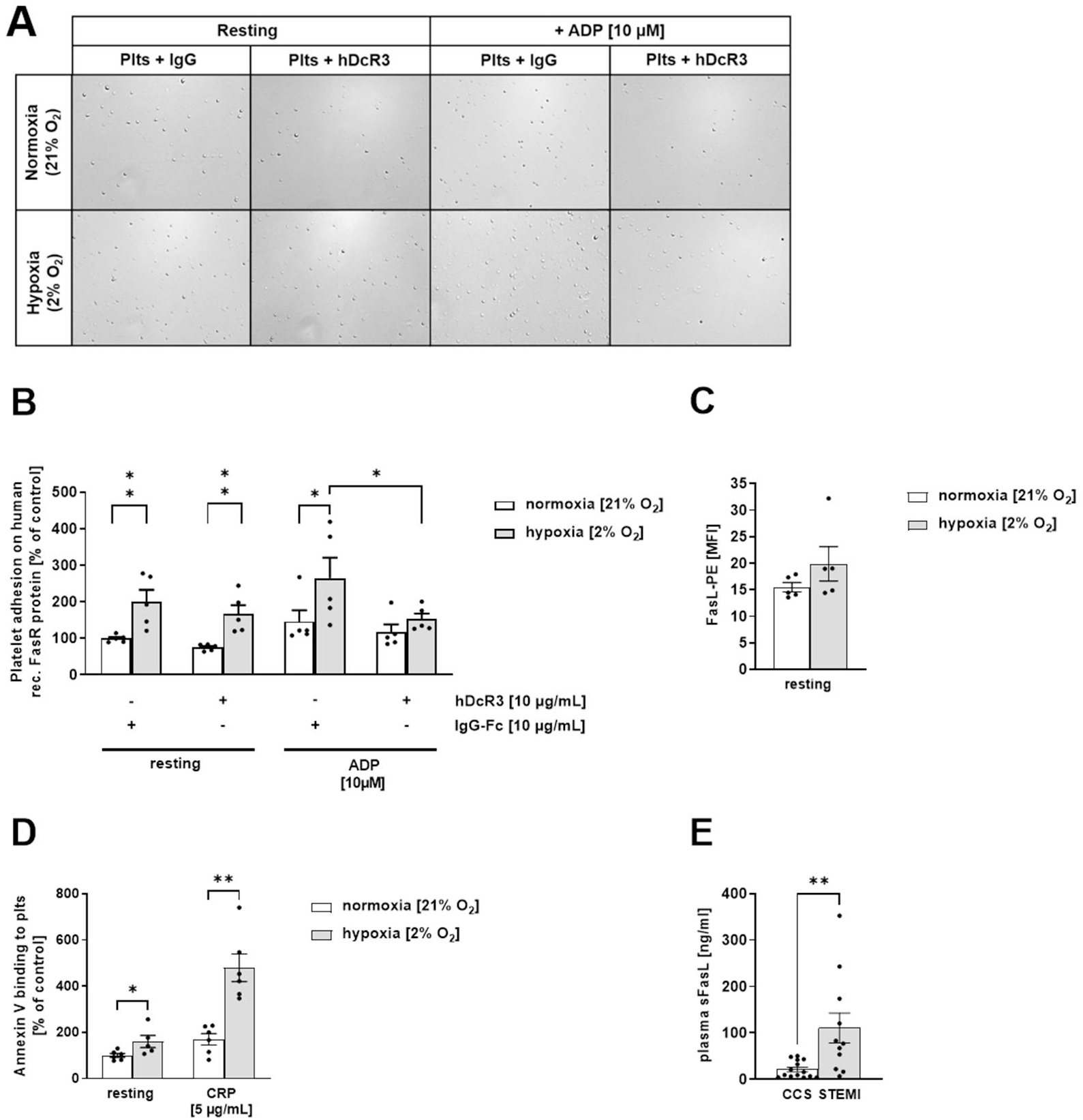
Increased adhesion of platelets to immobilized FasR under hypoxic conditions. **(A)** Representative images and (**B)** Quantification of adherent resting and ADP [10 μM]-stimulated human platelets to immobilized human recombinant FasR protein (50 μg/mL) under normoxic (21% O_2_) or hypoxic (2% O_2_) conditions (n = 4–6). (**C)** Externalization of FasL on the surface human platelet determined by flow cytometry (n = 5). (**D)** Annexin V binding to resting and CRP [5 μg/mL]-stimulated human platelets under normoxic (21% O_2_) or hypoxic (2% O_2_) conditions (n = 5-6) determined by flow cytometry. (**E)** Soluble (s)FasL plasma level (in ng/mL) from patients with coronary heart disease (CHD) compared to ST-elevation myocardial infarction (STEMI)-patients (n = 11–14). Scale bar = 50 μm (missing). Data are presented as means ± SEM. Statistical analyses were done by **(B)** Two-Way ANOVA followed by Sidak post hoc; **(C and E)** by two-tailed unpaired Student’s t-test and **(D)** multiple t-test. *p < 0.05, **p < 0.01, ***p < 0.001.

Platelet activation with ADP where the FasL was blocked by hDcR3 resulted reduced platelet adhesion compared to IgG-Fc controls (Fig. 3A-B) suggesting that the increase in platelet adhesion upon stimulation with ADP under hypoxic conditions is mediated by FasL binding of platelets to recombinant FasR. FasL exposure of platelets was enhanced by trend under hypoxic condition (Fig. 3C) while Annexin-V binding was significantly enhanced under hypoxia as shown by the analysis of resting and CRP stimulated platelets (Fig. 3D). Further evidence for FasL to be involved in I/R injury after AMI was provided by data from STEMI patients showing elevated plasma levels of sFasL compared to the CCS control group (Fig. 3E).

Consequently, we analyzed FasL exposure at the surface of platelets after 6h and 24h of ischemia and reperfusion (Fig. 4A). The exposure of FasL at the platelet membrane was significantly reduced 6h and 24h after I/R injury in wildtype C57BL/6 mice. In line with reduced platelet FasL exposure, we detected elevated plasma levels of sFasL in these mice 24h and 5 days after AMI suggesting that I/R injury leads to enhanced shedding of FasL after AMI (Fig. 4B).

**Figure 4.**
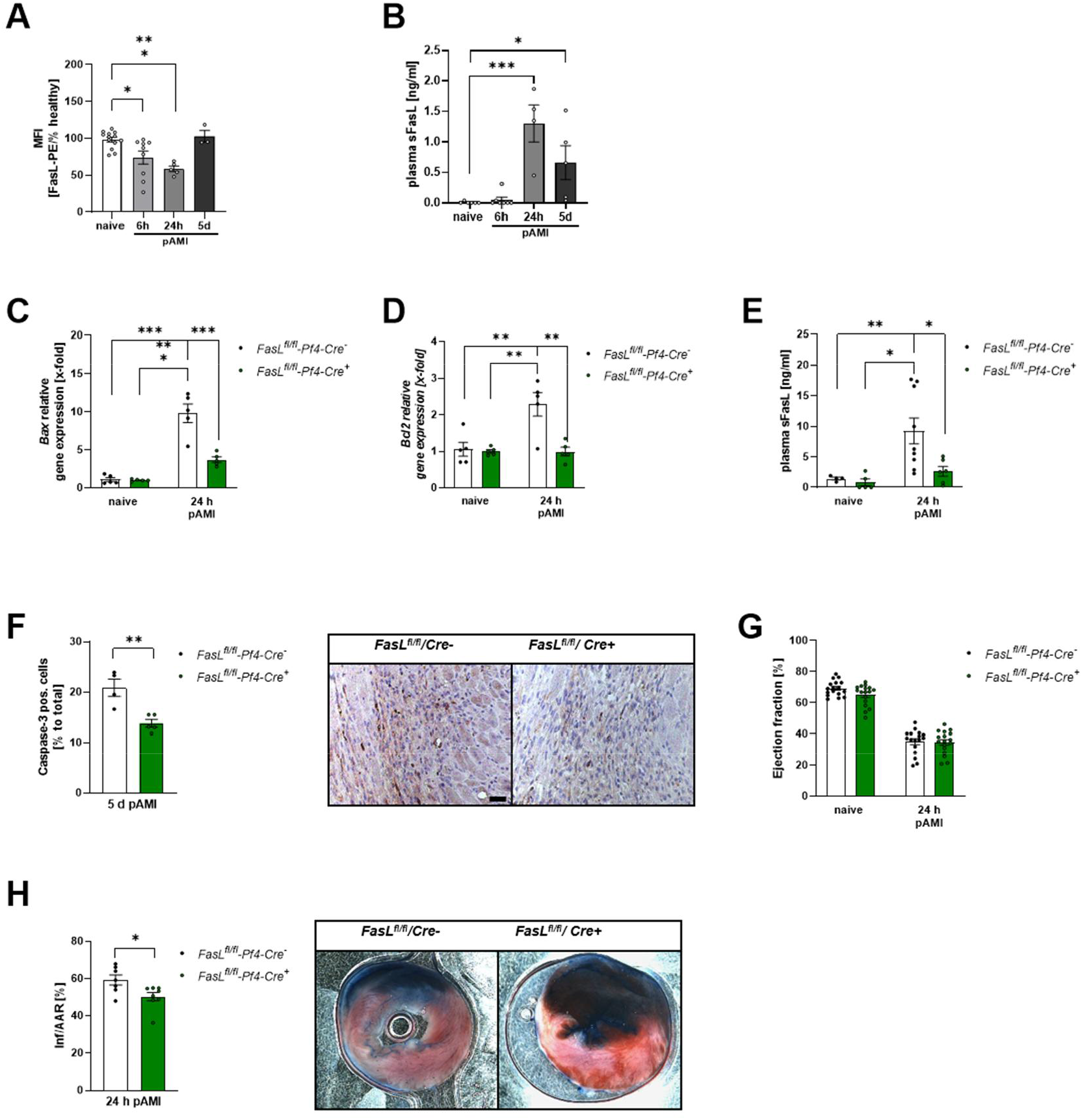
Genetic deficiency of platelet-specific FasL leads to reduced apoptosis post AMI. (**A**) FasL exposure at the platelet surface of naive mice and and 6h, 24h and 5 days post AMI; n = 13. (**B**) Plasma levels of soluble FasL (sFasL) in naive mice and 6h, 24h and 5 days post AMI; n = 5-7. (**C-D**) mRNA analysis of pro-apoptotic *Bax* (**C**) and anti-apoptotic *Bcl-2* (**D**) genes in the LV of *FasL*^*fl/fl*^*-Pf4-Cre*- and *FasL*^*fl/fl*^*-Pf4-Cre*+ naive mice and 24h post AMI; n = 5. (**E**) Plasma levels of sFasL in *FasL*^*fl/fl*^*-Pf4-Cre*- and *FasL*^*fl/fl*^*-Pf4-Cre*+ naive mice and 24 h post AMI; n = 4-9. (**F**) Quantification (left panel) and representative images (right panel) of caspase-3 positive cells in the heart of *FasL*^*fl/fl*^*-Pf4-Cre*- and *FasL*^*fl/fl*^*-Pf4-Cre*+ mice 5 days after AMI; n = 4-5. (**G**) Echocardiographic analysis of cardiac function by determination of ejection fraction (baseline vs. 24h after I/R) of *FasL*^*fl/fl*^*-Pf4-Cre*- and *FasL*^*fl/fl*^*-Pf4-Cre*+ mice; n = 12-13. (**H**) Quantitative analysis (left panel) of infarct size as the percentage of the area at risk (% Inf/AAR) with representative images (right panel) of *FasL*^*fl/fl*^*-Pf4-Cre*- and *FasL*^*fl/fl*^*-Pf4-Cre*+ mice 24 h post AMI. Blue = healthy tissue, red = the area at risk (AAR), white = infarcted area (INF); n = 7. Scale bar = 50 μm. Data are presented as means ± SEM. Statistical analyses were done by Two-Way ANOVA followed by Sidak post hoc **(A-E, G)** or by two-tailed unpaired Student’s t-test **(F and H)**. *p < 0.05, **p < 0.01, ***p < 0.001.

### 3.4 Platelet FasL exposure mediates cell apoptosis in the left ventricle post AMI

According to the results shown in figure 3, we analyzed the impact of platelet-specific loss of FasL on cell apoptosis post AMI using *FasL*^*fl/fl*^*-Pf4-Cre+* mice that do not express FasL on the platelet surface (26). *FasL*^*fl/fl*^*-Pf4-Cre*mice were used as wildtype controls. Accordingly, AMI induced significantly elevated *Bax* gene expression in the LV of *FasL*^*fl/fl*^*-Pf4-Cre*mice after 24h of reperfusion compared to naïve conditions (Fig. 4A). However, *FasL*^*fl/fl*^*-Pf4-Cre+* mice showed only a weak increase of LV Bax expression compared to naive conditions and was significantly lower compared to *FasL*^*fl/fl*^*-Pf4-Cre-*. Further, AMI also induced LV *Bcl-2* expression in FasL wildtype mice (Fig. 4B). Similarly, to *Bax* expression, *FasL*^*fl/fl*^*-Pf4-Cre*+ mice showed almost no upregulation of *Bcl-2* expression after AMI compared to naïve mice and less *Bcl-2* expression compared to *FasL*^*fl/fl*^*-Pf4-Cre-* (Fig. 4B). In addition, the analysis of plasma levels of sFasL in *FasL*^*fl/fl*^*-Pf4-Cre-* and *FasL*^*fl/fl*^*-Pf4-Cre*+ post AMI revealed a significant increase in *FasL*^*fl/fl*^*-Pf4-Cre*-mice that was strongly reduced in *FasL*^*fl/fl*^*-Pf4-Cre*+ mice 24h after AMI, suggesting that platelets are major contributors of sFasL plasma levels after AMI (Fig. 4C). Furthermore, histological analyses revealed that the loss of platelet FasL is responsible for reduced caspase-3 activity in the LV of *FasL*^*fl/fl*^*-Pf4-Cre*+ mice after 5 days of reperfusion (Fig. 4D). Nevertheless, the FasL-associated changes in LV apoptosis alone had no influence on heart function post AMI (Fig. 4E). However, after 24h of reperfusion, we detected a reduced infarct size in *FasL*^*fl/fl*^*-Pf4-Cre*+ mice compared to controls (Fig. 4F) due to reduced cell apoptosis after ischemia and reperfusion injury.

**Table 1.**
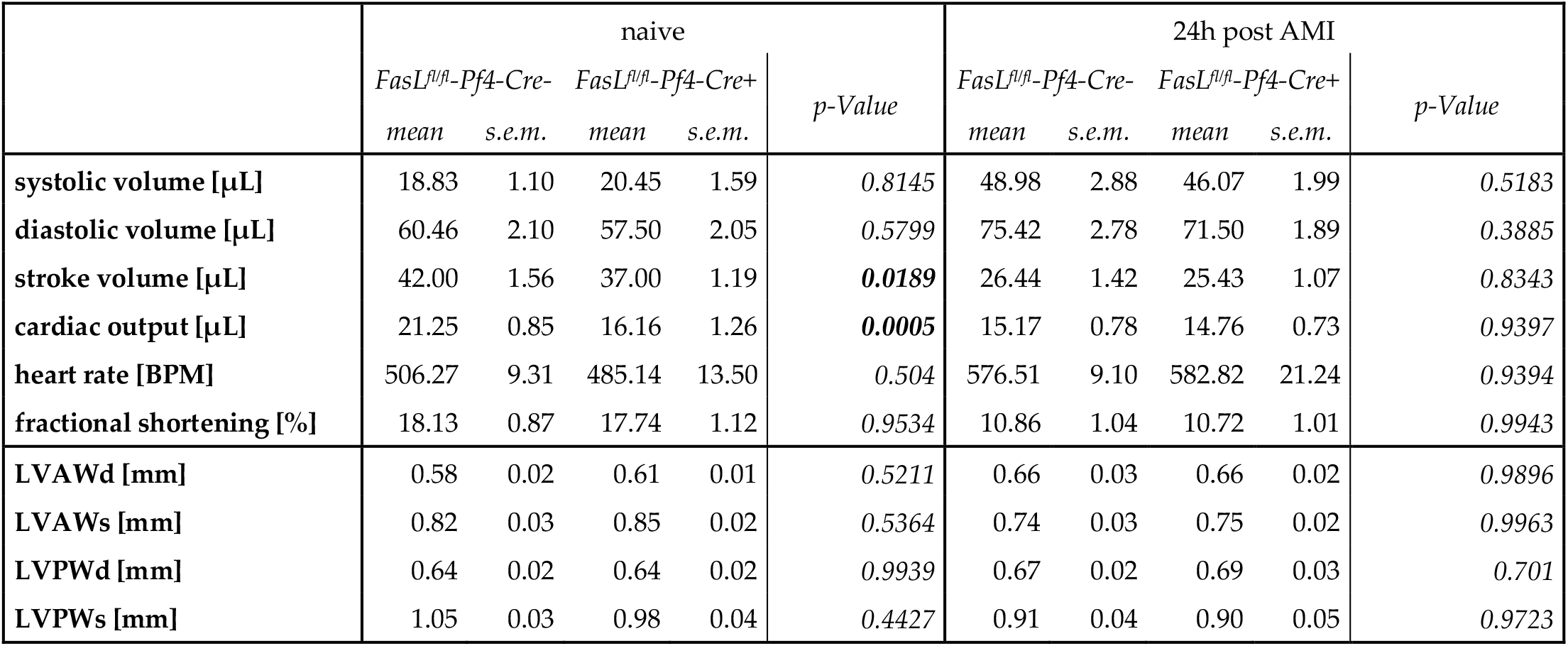
Different parameters of LV function and wall thickness of mice after AMI. Echocardiographic analysis was conducted to evaluate heart function and wall thickness parameters in *FasL*^*fl/fl*^*-Pf4-Cre* mice following ischemia/reperfusion (I/R) injury (n=17). Statistical significance was determined using Two-Way ANOVA with Sidak’s multiple comparison test. Data are presented as mean ± SEM. LVAWs = left ventricular anterior wall thickness during systole; LVAWd = left ventricular anterior wall thickness during diastole; LVPWs = left ventricular posterior wall thickness during systole; LVPWd = left ventricular posterior wall thickness during diastole.

## 4. Discussion

In this study, we found that I/R injury leads to apoptosis of resident cardiac cells after AMI in mice. Platelets are one major contributor of cell apoptosis because thrombocytopenia and genetically modified mice with a loss of FasL restricted to platelets resulted in reduced cardiac cell death via the intrinsic and the extrinsic pathway of apoptosis. Mechanistically, we provide first evidence that platelet binding to FasR is significantly elevated under hypoxic conditions as found in AMI.

In AMI, apoptosis is a key pathological determinant since the inhibition of apoptosis has a beneficial effect in experimental studies. Thus, to understand the mechanisms that lead to apoptosis is important for therapeutic options to prevent and/or treat heart failure induced by ischemic reperfusion injury but also for diagnosis and risk stratification in patients with ischemic heart disease (27). To date, it is still not completely understood to which extent apoptosis counts for cardiac damage because of the large differences in reports from different groups. There might be also differences in the induction of apoptosis by ischemia and/or by reperfusion. Ischemia alone did not result in DNA laddering, a marker of apoptosis, but was observed after reperfusion indicating that apoptosis in the myocardium is triggered upon reperfusion but does not manifest during the ischemic event. In contrast, others found that myocardial ischemia induces apoptosis when a prolonged ischemic period without any reperfusion was observed while others detected apoptosis after brief ischemic events followed by reperfusion (4, 28). However, different studies provided evidence for the initiation of apoptosis upon ischemia as indicated by pro-apoptotic markers and caspase activation but no DNA fragmentation followed by a massive increase during reperfusion. Thus, it becomes evident that apoptosis might be fully conducted during reperfusion. The assumption that reperfusion is a trigger for apoptosis that has been further strengthened by a study where inhibitors of pro-apoptotic mediators were used because this resulted in reduced infarct size at early reperfusion (6). Studies in humans provided evidence for long-lasting cell apoptosis in the myocardium because they detected apoptotic cardiomyocytes in the border zone of the infarcted myocardium within hours to days of infarction (29). On the contrary, minimal expression of proteins of the apoptosis signaling cascade in cardiomyocytes argues against a significant role of apoptosis in I/R induced cell death (30). This hypothesis has been reinforced by knock-out mice with cardiac specific ablation of caspase-3 and caspase-7 that do not show beneficial effects for infarct size or LV remodeling indicating no acute effects on myocardial I/R injury (31). Thus, it is not surprising that overexpression of caspase in cardiomyocytes does not lead to increased apoptosis in cardiomyocytes but elevated infarct size during I/R (32). These results together implicate that the previous observations of apoptosis in the heart are likely to have been due to apoptosis of non-cardiomyocytes (30). This hypothesis is emphasized by the idea that apoptosis of endothelial cells and leucocytes might indirectly affect cardiomyocyte cell survival and cardiac performance (30). This suggests that anti-apoptotic strategies might be still cardio-protective, particular with regard to a recently published study with a new developed peptide targeting the Fas-dependent apoptotic signal during IR injury (33). In line with our study using platelet-specific FasL knock-out mice, decreased infarcts size was shown when the Fas-dependent apoptotic pathway was blocked with this peptide, suggesting that the FasL-FasR signaling pathway is a promising target to reduce myocardial injury after AMI. Importantly, hypoxia plays a role in FasL-FasR induced cell apoptosis because reduced oxygen supply triggers increased expression of FasR on cardiomyocytes on the one hand (34, 35) and the ability of platelets to adhere to FasR on the other hand (Figure 3).

Platelets play a major in ischemic cardiovascular disease. Beyond thrombotic complications, they modulate inflammation and cardiac remodeling (13, 14). Here, we provide strong evidence for platelets to induce cell apoptosis of resident cardiac cells via exposure of FasL. Data from Mpl deficient and platelet depleted mice clearly show a significant contribution of platelets to cell death via apoptosis induced by the intrinsic and the extrinsic pathway. Recently, we have shown that platelets are the major source of IL-1β after I/R because IL-1β plasma levels were almost absent in platelet depleted mice 24h post AMI (14). However, IL-1β is not only a crucial acute phase cytokine important to induce the activation and secretion of IL-1β from leukocytes but is also responsible for the increase in FasL-induced caspase-3 activation as shown in hepatocytes (36). Thus, reduced apoptosis in platelet depleted mice after AMI might be provoked by different mechanisms such as reduced IL-1β plasma levels, reduced hypoxia and absent platelet-FasL induced cell apoptosis of migrated platelets in the infarct zone. Beside reduced intrinsic, mitochondrial cell death as a result of reduced hypoxia in platelet depleted mice, we provided evidence that platelet FasL induces cell death in the infarct zone of the LV. Platelet-specific genetic deficiency of FasL led to reduced apoptosis in the LV and thus affects infarct size but not cardiac function. In the past, platelet FasL has been shown to trigger cell death in murine neuronal cells and in a murine stroke model (18). It was shown that *Bax/Bak*-induced mitochondrial apoptosis signaling in target cells was not required for platelet-induced cell death but increased the apoptotic response to platelet-FasL-induced Fas signaling. Here, we also found reduced *Bax* and *Bcl-2* expression in the LV of mice with platelet-specific FasL ablation (Fig. 4A-B). Thus, mitochondrial-induced cell death after AMI might trigger the observed platelet FasL-induced cell apoptosis in the LV. However, there might be additional mechanisms that account for reduced cell apoptosis in platelet depleted mice after AMI because additional death receptor ligands are known to be stored and secreted by platelets such as TRAIL (Apo2-L), TWEAK (Apo3-L) and LIGHT which have the capacity to regulate apoptosis via paracrine signaling (37-39).

Mitochondria play an important role in cell apoptosis because they are the major contributors of reactive oxygen species (ROS) and the major target for ROS-induced damage. Typical characteristics of the mitochondrial apoptosis pathway are mitochondrial swelling and outer mitochondrial membrane rupture that triggers the release of pro-apoptotic factors such as cytochrome c and SMAC/Diablo from the intermembrane space into the cytosol (40). Interestingly, chronic thrombocytopenia does not reduce mitochondrial (intrinsic) apoptosis in the myocardium of MPL knock-out mice as shown by unaltered expression of *Bax* and *Bcl*-*2*. As already shown by Schleicher and colleagues, mitochondrial apoptosis signaling is not required for platelet induced cell death (18). Thus, reduction of caspase-3 positive cells in the myocardium of MPL knock-out mice is the result of extrinsic apoptotic mechanisms probably via FasL mediated activation of FasR. However, different results of mice with chronic and acute thrombocytopenia might be also due to different platelet counts in these mice. While platelet depleted mice represent a platelet count of <01%, we detected ∼10% of platelet numbers in Mpl deficient mice compared to controls (14). Thus, it is tempting to speculate that differences around 10% of platelet counts might be sufficient for alterations in mitochondria mediated apoptosis in the myocardium.

Interestingly, the soluble form of FasL is not able to trigger apoptosis (41). After AMI, the exposure of FasL at the platelet surface was reduced at 6h and 24 h post I/R that correlates with enhanced plasma levels of FasL after AMI. However, our results clearly indicate that the majority of plasma sFasL is of platelet origin because plasma levels of sFasL are strongly reduced in platelet specific FasL knock-out mice (*FasL*^*fl/fl-*^*Pf4-Cre*+). Consequently, platelets might be involved in the prevention of cell apoptosis of non-resident cells in the circulation such as leukocytes by activation induced shedding of the FasL from the platelet surface.

## 5. Conclusions

Recent findings together with the here presented results of platelet induced cell apoptosis clearly indicate an important role of platelets for cardiac function beyond occlusive ischemic damage post AMI. Platelets support the acute inflammatory response and migrate into the infarct border zone to actively modulate cardiac remodeling and cell apoptosis. These new identified mechanisms of platelet induced cardiac damage needs to be taken into account while optimizing antithrombotic/antiplatelet strategies for patients with AMI.

## Supplementary Materials

None.

## Author Contributions

Conceptualization, M. E.; methodology, S.G.; validation, M.K., J.W.F., M.E.; investigation, K.J.K. F.R., M.D.,A.P.; resources, S.K.; writing—original draft preparation, M.E., F.R., K.J.K.; writing—review and editing, M.E.; funding acquisition, M.E. All authors have read and agreed to the published version of the manuscript.

## Institutional Review Board Statement

Animal studies were performed in accordance with the guidelines of the European Parliament for the use of living animals in scientific studies and in accordance with the German law for protection of animals. The protocol was approved by Heinrich-Heine-University Animal Care Committee and by the district government of North-Rhine-Westphalia (LANUV; NRW; Permit Number 84-02.04.2015.A558; 81-02.04.2019.A270; 84-02.04.2017.A440).

Experiments with human blood were reviewed and approved by the Ethics Committee of the Heinrich-Heine-University. Subjects provided informed consent prior to their participation in the study (patients’ consent). The experiments conform to the principles outlined in the Declaration of Helsinki.

## Informed Consent Statement

Informed consent was obtained from all subjects involved in the study.

## Data Availability Statement

The data presented in this study are available on request from the corresponding author.

## Acknowledgements

The authors would like to thank Martina Spelleken, Elena Schickentanz-Dey and Meike Klier for excellent performance of experiments. We would like to acknowledge the Center for Advanced Imaging (CAi) at Heinrich-Heine-University Düsseldorf for providing access to the confocal microscope from Zeiss (LSM 880 Airyscan).

## Disclaimer/Publisher’s Note

The statements, opinions and data contained in all publications are solely those of the individual author(s) and contributor(s) and not of MDPI and/or the editor(s). MDPI and/or the editor(s) disclaim responsibility for any injury to people or property resulting from any ideas, methods, instructions or products referred to in the content.

